# Emergence of Probabilistic Representation in the Neural Network of Primary Visual Cortex

**DOI:** 10.1101/2021.02.06.429971

**Authors:** Ang A. Li, Fengchao Wang, Xiaohui Zhang, Si Wu

## Abstract

During the early development of mammalian visual system, the distribution of neuronal preferred orientations in the primary visual cortex (V1) gradually shifts to match the major orientation features of an environment, achieving optimal representation of the environment. By combining the computational modeling and experimental electrophysiological recording, we provide a circuitry plasticity mechanism that underlies the developmental emergence of such matched representation in the visual cortical network. Specifically, in a canonical circuit of densely interconnected pyramidal cells and inhibitory parvalbumin-expressing (PV+) fast-spiking interneurons in the V1 layer 2/3, our model successfully simulates the experimental observations and further reveals that the non-uniform inhibition, mediated by local interneurons, exerts a key role in shaping the network representation through spike timing-dependent synaptic modifications. The experimental results confirm that PV+ interneurons in the V1 are capable of providing such non-uniform inhibition during a short period after the vision onset. Thus, our study elucidates a circuitry mechanism for acquisition of the prior knowledge of environment for optimal inference in sensory neural system.

## Introduction

Neural networks of animal and human develop internal representations that match the environment and guide behaviors. In the mammalian primary visual cortex (V1), neurons encode oriented bars or edges of light in their receptive fields (***Hubel and Wiesel, 1959***). Collectively, these neural population can represent the statistics of the visual environment (***Pouget et al., 2000***). Recent framework for probabilistic representation in sensory cortex propose that the prior distribution of stimuli can be represented by the distribution of preferred orientation of the neural population (***Fischer and Peña, 2011*; *Girshick et al., 2011*; *Ganguli and Simoncelli, 2014***). Indeed, such proposal is supported by the experimental observations: under normal condition, neurons in the V1 of adult animals and human show an over-representation of cardinal orientations, which coincides with the local orientation distribution measured in photography (***Chapman and Bonhoeffer, 1998*; *Furmanski and Engel, 2000*; *Girshick et al., 2011*; *Hagihara et al., 2015***); while the altered environment - the restriction of visual experience to contours of only one orientation (stripe rearing) - leads to an over-representation of the experienced orientations (***Stryker et al., 1978*; *Kreile et al., 2011***). However, how such probabilistic representation emerges in the development of cortical network is not known (***Seriès and Seitz, 2013***).

We aim to address this question by examining the development of a canonical microcircuit that has been proposed as the building block for predictive coding (***Bastos et al., 2012***) and Bayesian inference (***Darlington et al., 2018*; *Nessler et al., 2013***). The microcircuit consists densely interconnected populations of excitatory and inhibitory neurons (***Hofer et al., 2011***): previous experiments suggested that the connectivity between PCs and inhibitory PV+ interneuron in the cortical layer 2/3 are random, reciprocal and dense -with connection probability reaching 50% (***Miao et al., 2016*; *Avermann et al., 2012***). This microcircuit is largely matured early before the vision onset (or eye-opening,EO) and shapes the entire process of visual development (***Magueresse and Monyer, 2013***), including the sharpening of orientation tuning and the onset of a critical period (CP) of experience-dependent neuronal connection refinements during the early postnatal development (***Lee et al., 2012*; *Kuhlman et al., 2013***).

In computational studies, this microcircuit is frequently modeled as a feedback inhibition network (***Avermann et al., 2012***). Combined with a synaptic learning rule of spike timing-dependent plasticity (STDP), it has been used to study the development of V1, including emergence and sharpening of orientation tuning (***Song and Abbott, 2001*; *Sadeh et al., 2015***). We propose that this feedback inhibition network is ideal for the probabilistic representation: Different from winner-take-all (WTA) inhibition (***Rumelhart and Zipser, 1985***), feedback inhibition provides a softer form of inhibition in the cortical network (***Avermann et al., 2012*; *Jonke et al., 2017***), where the level of inhibition is proportional to the level of excitation (***Isaacson and Scanziani, 2011***). This softer form of inhibition allows a subset of excitatory neurons to respond to a specific input feature, which enables the neural population to represent a probability distribution.

We combine mathematical modeling with electrophysiology experiment to study the emergence of probabilistic representation in this cortical microcircuit, with emphasize on the role of PV+ inhibitory neurons - the major player of feedback inhibition. Based on the feedback inhibition network model with STDP, we build a computational model that mimics the development of cortical microcircuit under normal and altered environments. Complementarily, we perform *in vivo* electrophysiology recordings to measure the tuning properties of PCs and inhibitory PV+ interneurons in the mouse V1 across different developmental stages: shortly after the EO, during the CP, and in the adulthood. Our modelling and experimental results show a tight relationship between the distributions of preferred orientations of PCs and inhibitory PV+ interneurons across different development stages, and delineate a crucial role of the feedback inhibition in the developmental emergence of probabilistic representation in the cortical network.

## Results

### Emergent properties of the feedback inhibition network

We study the emergence of probabilistic representation in the cortical network using a ubiquitous microcircuit model in the brain, referred as the feedback inhibition network (***Grossberg, 1976*; *Jonke et al., 2017***). The network mimics the densely interconnected excitatory and inhibitory neurons in the cortex. Previous experiments showed that the connectivity between PCs and PV+ interneurons in layer 2/3 is random, reciprocal, and dense - with the connection probability reaching 50% (***Miao et al., 2016*; *Avermann et al., 2012***). This microcircuit matures early before EO and shapes the entire process of visual development including the sharpening of orientation tuning and the onset of CP (***Lee et al., 2012*; *Magueresse and Monyer, 2013*; *Kuhlman et al., 2013***). Moreover, this microcircuit has been proposed as the building block for predictive coding (***Bastos et al., 2012***) and Bayesian inference (***Darlington et al., 2018*; *Nessler et al., 2013***)..

The model consists of interconnected excitatory and inhibitory neural populations, of which 80% are excitatory neurons and 20% inhibitory (***Figure 1***A). Inhibitory neurons are recurrently connected. The connection probabilities between different type of neurons are in accordance to the experiment data of visual cortex (***Miao et al., 2016***). Each neuron is modeled as leaky integrate-fire neuron.

**Figure 1.**
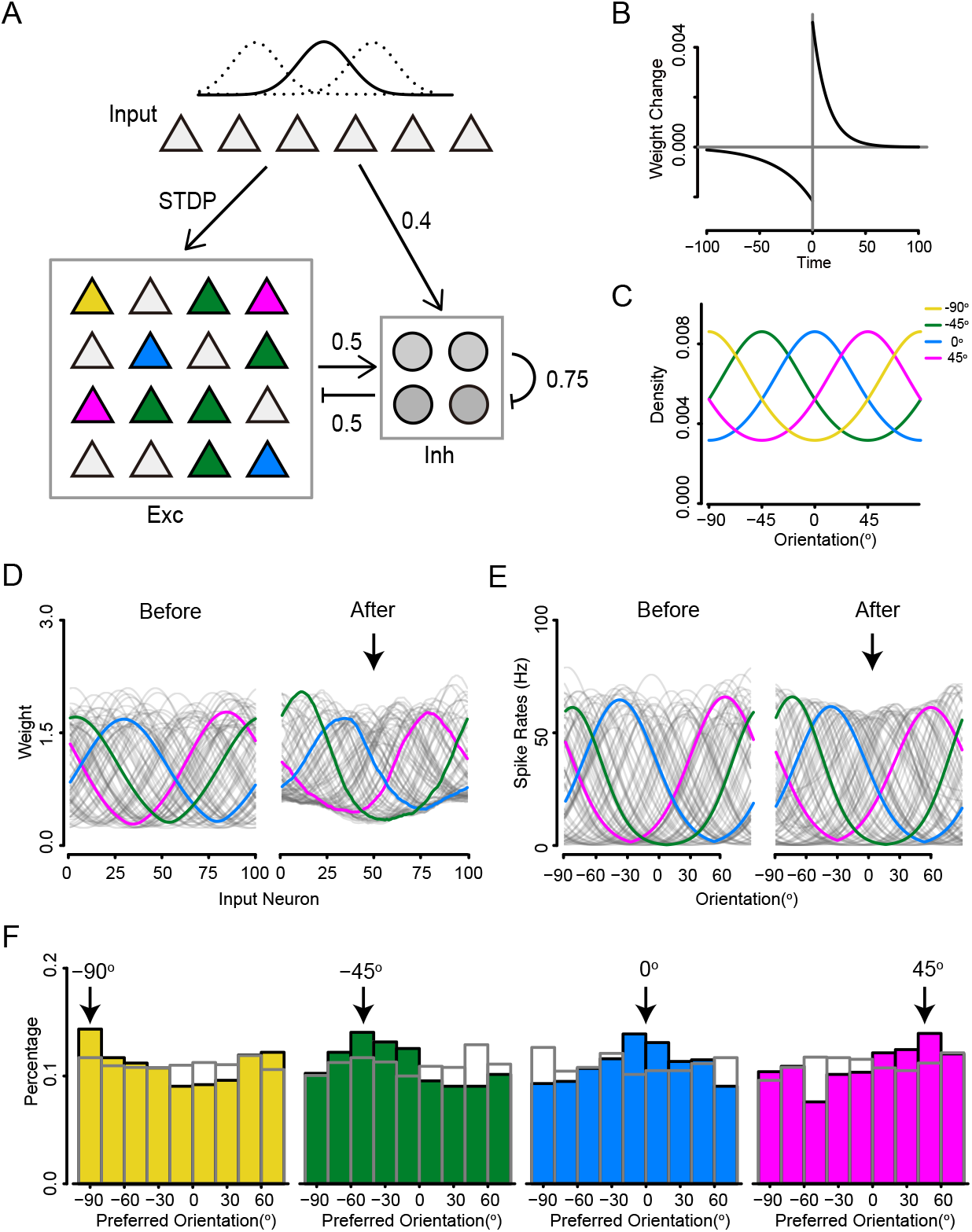
Emergent properties of the cortical network under stripe rearing condition. **A**. Schematic diagram of the network model structure. Input neurons encodes an orientation variable with a Gaussian profile of their firing rates. Triangles represents excitatory population and circles represents inhibitory population. Colors represent neurons with different preferred stimulus. **B**. The standard STDP learning rule used in the model, which specifies the change of synaptic weight for a pre-post paring with the time difference Δ*t* = *t*_*post*_ − *t*_*pre*_ between a postsynaptic spike at time *t*_*post*_ and a presynaptic spike at time *t*_*pre*_. **C**. Orientation input encoded by 100 Poisson neurons with a Gaussian profile of firing rates. **C**. Prior probabilities of orientations under different stripe rearing conditions. Yellow, green, blue and magenta lines indicate experienced orientation at -90°, -45°, 0°, and 45°, respectively. **D**. feed-forward connection weights of simulated excitatory neurons before and after training under stripe rearing condition at 0 ° (smoothed by a moving-average filter with 10 nearest points). Three sample neurons are highlighted with magenta, green, and blue, respectively. **E**. Simulated excitatory neurons before and after training under stripe rearing condition at 0 °. The same neurons in **D**. are highlighted with magenta, green and blue, respectively. **F**. Histograms of preferred orientations under each rearing condition, experienced orientations are specified above the graphs. Grey bars represent initial conditions.

Both excitatory and inhibitory neurons receive feed-forward inputs modeled as Poisson spikes. But only the feed-forward connections received by excitatory neurons are plastic and subject to a standard STDP rule (***Figure 1***B), with homeostatic weight rescaling. Similar models have been used to study the visual development (***Song and Abbott, 2001*; *Sadeh et al., 2015*; *Jonke et al., 2017***), but how the probabilistic representation emerges in such type of networks is not well understood.

We examine the emergent properties of the feedback inhibition network in a setting that mimics the stripe rearing condition: the restriction of visual experience to contours of only one orientation. Previous experiments showed that stripe rearing leads to an over-representation of the experienced orientation among neurons in the visual cortex (***Kreile et al., 2011*; *Stryker et al., 1978***). In our simulation, the orientation input is encoded by Poisson spiking neurons with a Gaussian profile of firing rates (***Song and Abbott, 2001*; *Sadeh et al., 2015***). Before training, the excitatory neurons are initialized to be orientation selective, with uniformly distributed preferred orientations (***Figure 1***F, gray bar). To mimic the stripe rearing condition, the prior distribution of orientation inputs follows a von-Mises distribution that peaks at the experienced orientation (***Figure 1***C). We select 4 stripe rearing conditions with experienced orientations at -90°, -45°, 0°, 45°, respectively (***Kreile et al., 2011***).

Stripe rearing has a strong effect on simulated neurons. After training, the feed-forward connection weights of simulated neurons tend to shift towards the experienced orientation (***Figure 1***D). As a result, the tuning curves also shift towards the experienced orientation (***Figure 1***E). On the population level, the distribution of preferred orientation shows an over-representation at the experienced orientation (***Figure 1***F), which is consistent with the experiment observation. To quantify the specific effect due to stripe rearing, we compare the increase in the fraction of neurons preferring the experienced orientation to the decrease in preference for the orthogonal orientation (***Kreile et al., 2011***). The average specific effect for stripe rearing is 12.68% (-90°: 10.65 ± 8.55%, -45°: 13.95 ± 8.01%, 0°: 13.05 ± 5.13% and 45°: 13.05 ± 6.39%). Our results suggest that the distribution of preferred stimuli in the feedback inhibition network will shift according to the prior distribution of the input, leading to the stripe rearing effect.

### Emergence of the overrepresentation of cardinal orientations in the cortical network

During visual development, neurons in V1 develop an overrepresentation of cardinal orientations (***Hagihara et al., 2015***), which coincides with the local orientation distribution measured in photography (***Girshick et al., 2011***) (***Figure 2***A). This process can be understood as the neural network matching its internal model to the environment for optimal inference (***Girshick et al., 2011*; *Fischer and Peña, 2011*; *Ganguli and Simoncelli, 2014***). Here, we combine computational and experimental methods to study its circuit mechanism.

**Figure 2.**
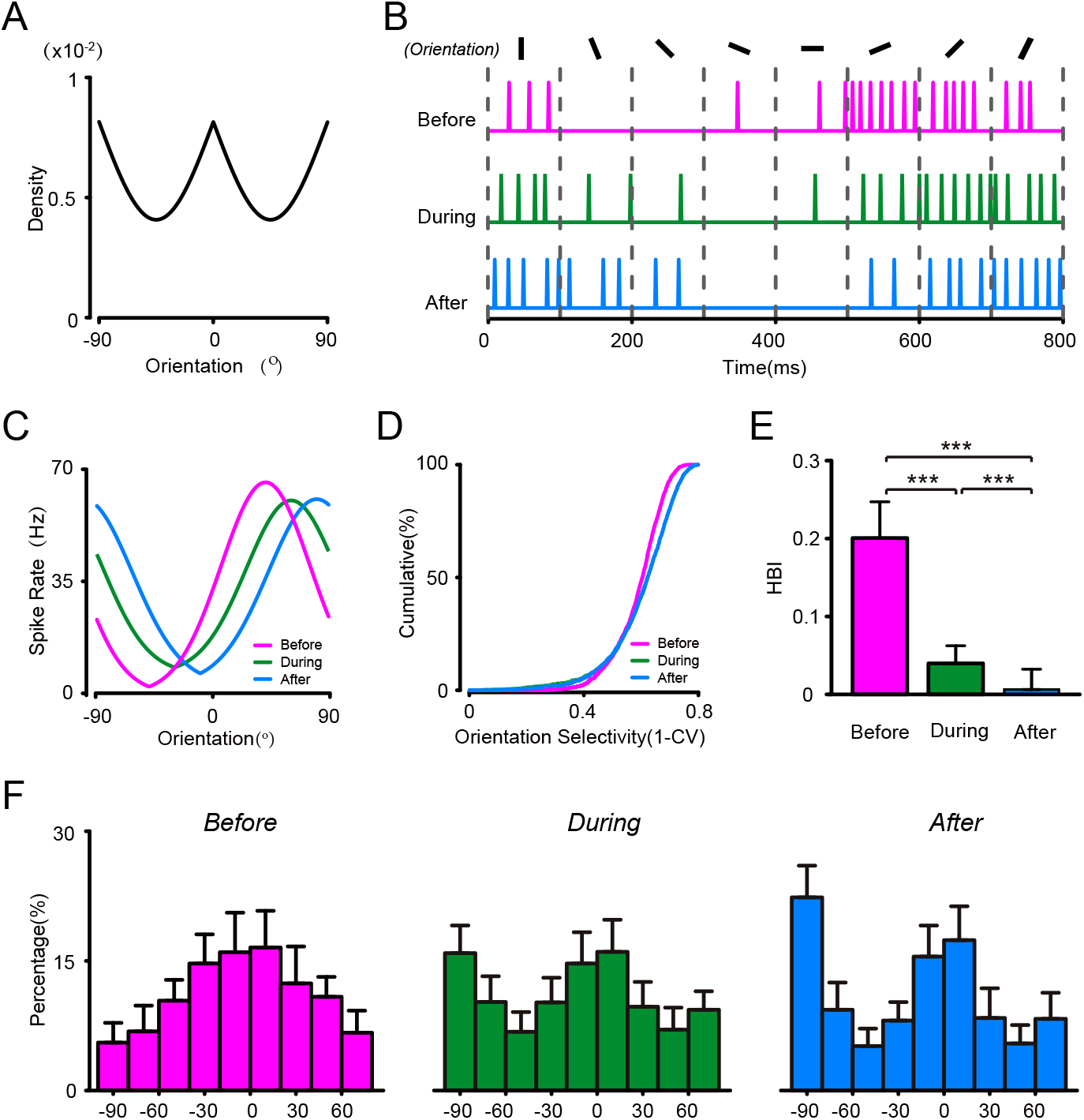
Emergence of the overrepresentation of cardinal orientations in the feedback inhibition model **A**. Prior probabilities of the orientations in natural images. **B**. Spike trains of a representative simulated excitatory neuron to stimuli with 8 different orientations, before (magenta), during (green) and after (blue) learning. **C**. Orientation tuning curves of a simulated excitatory neuron at different learning stages (the same neuron as in **B**). **D**. Distribution of orientation selective index (OSI) of simulated excitatory neurons at different learning stages. **E**. Horizontal bias index (HBI) of simulated excitatory neurons at different learning stages. **F**. Distribution of preferred orientation of simulated excitatory neurons before (left), during (middle), and after (right) learning. P values in **E**. are calculated by the Kolmogorov–Smirnov test. *** indicates P<0.01; ** indicates 0.01<P<0.05.

We study the emergent properties of the feedback inhibition network under a condition that mimics visual development. Experiments find that, for mice few days after EO, the distribution of preferred orientation in PCs shows a horizontal bias, with more neurons preferring horizontal orientations (***Hagihara et al., 2015***). Thus, we initialize the preferred orientations of excitatory population with a von-Mises distribution that peaks at horizontal orientation (***Figure 2***F left). During learning, orientation inputs are randomly drawn from a prior distribution, mimicking the local orientations measured in natural images (***Girshick et al., 2011*; *Wei and Stocker, 2015***) (***Figure 2***A). The tuning properties of neurons are tested by 8 different orientation stimuli before, during and after the learning period (***Figure 2***B).

We find the over-representation of cardinal orientation emerges in the network model. During learning, preferred orientation of excitatory neurons tend to shift towards vertical orientation, which is initially under-representated by the network. On the population level, the horizontal bias in the distribution of preferred orientation of excitatory neurons becomes cardinal, with peaks at both horizontal and vertical orientations (***Figure 2***F), which is stable during later half of the training period. To quantify the changes, we quantify the horizontal bias using horizontal bias index (HBI) (***Hagihara et al., 2015***). The excitatory neurons before learning has a significantly larger HBI than those during and after learning (***Figure 2***E, t-test between before and during learning, P < 2.2E-16; between before and after learning, P < 2.2E-16). However, the distribution of orientation selectivity index (1-CV) does not change significantly during training (***Figure 2***D), suggesting the over-representation of cardinal orientation is solely caused by the shift of preferred orientation of excitatory neurons. Our model successfully reproduces the emergence of the over-representation of cardinal orientation during visual development.

To validate our model results, we use *in vivo* extracellualr single-unit recording to measure the orientation selectivity of PCs in the binocular area of the mouse V1 at different development stages: shortly after the EO (postnatal day 17-18, P17–18), during the CP (P27-28) and in the adult stage (P55-56). For each developmental stage, we measure spiking activity of putative PCs in response to drifting grating stimuli with 12 different directions (6 orientations) (***Figure 3***A & B). We compute the preferred orientation of each PC by fitting the spike-rate vs orientation tuning curve with a Gaussian function, and calculate the orientation selective index (OSI) for measuring their orientation tuning amplitudes (***Figure 3***C). We find that for PCs recorded shortly after the EO, the OSI amplitudes have already had a comparable level as that during the CP (two-sample Kolmogorov– Smirnov test between P17-P18 and P27-28, p = 0.557) or the later adult stage (two-sample Kolmogorov– Smirnov test between P17-P18 and P55-56, p = 0.053) (***Figure 3***D), which confirms the previous findings that the orientation selectivity of PCs are largely matured before the EO (***Hagihara et al., 2015*; *Hoy and Niell, 2015***).

**Figure 3.**
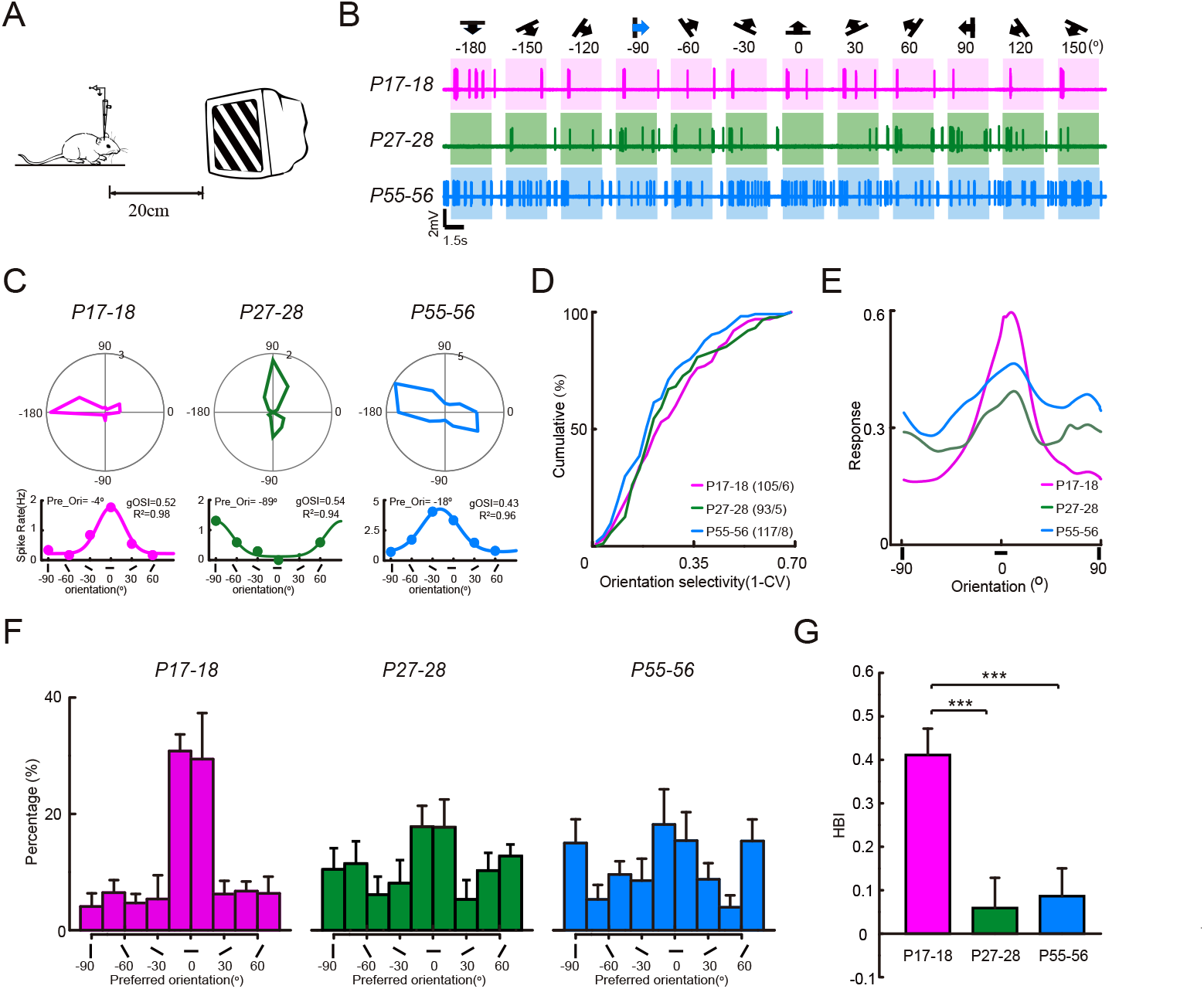
Emergence of the overrepresentation of cardinal orientations in the cortical network of V1 during development. **A**. Schematic diagram illustrating *in vivo* single-unit recording with glass micropipettes of spike responses from layer 2/3 PCs in mouse V1. **B**. The response of representative neurons in different postnatal stage to presented drifting gratings with 12 different directions (magenta trail, P17-18; green trail, P27-28; blue trail, P55-56). The stimulus directions and orientations are indicated by the symbols shown on top. **C**. Top, the direction-tuned responses of representative neurons in **A**. Bottom, the curves denote the method to use one-peak Gaussian function fitting to detect the preferred-orientation of representative neurons in **B. D**. Cumulative distributions of gOSI (1-CV) values of PCs at 3 different postnatal stages (magenta line, P17-18; green line, P27-28; blue line, P55-56). **E**. Population tuning curves of PCs at 3 different postnatal stages (magenta line, P17-18; green line, P27-28; blue line, P55-56). **F**. Distributions of preferred orientations of PCs in P17-18 mice (98 cells / 6 mice), P27-28 mice (89 cells / 5 mice) and P55-56 mice (113 cells / 8 mice). **G**. HBI of PCs in 3 different postnatal stages. P values are calculated by the Kolmogorov–Smirnov test and t-test in **D**. and **G**. respectively. *** indicates P<0.01; ** indicates 0.01<P<0.05.

At the population level, we analyze the distribution of preferred orientation of putative PCs at three developmental stage. The distribution shows an apparent bais to the horizontal orientation in those mice examined shortly after the EO, consistent with previous findings (***Hagihara et al., 2015***). However, During the CP the distribution evolves to be a bi-modal with two peaks at the horizontal and vertical orientations, which further remains stable through the adult stage (***Figure 3***F). The horizontal biases are agin quentified using HBI. We find that the PCs shortly after EO has a significantly stronger bias toward the horizontal orientations than those during the CP (t-test, p = 1.75E-04) and the adult stage (t-test, p = 3.22E-04, ***Figure 3***G). Moreover, we also compute the population tuning curve for the recorded PC population by averaging the normalized tuning curves of individual PCs (***McAdams and Maunsell, 1999***). The population tuning curve quantitatively reflect the population activity to different orientations. We observe the population activity of recorded PCs clearly shows a bias to the horizontal orientation shortly after the EO, while the latter bias become absent during the CP and in the adult (***Figure 3***E). Such developmental change in the population tuning curve of PC agrees with the observed developmental evolution of the neuronal preferred orientation distribution. Thus, these experimental recordings directly reveals a switching process from the over-representation of horizontal orientation to the cardinal orientations in the V1 layer 2/3 PCs during a period before the CP, and these results verify the model simulations.

### Inhibitory neurons provide non-uniform feedback inhibition during early development stage

Inhibitory neurons play a key role in the visual development (***Espinosa and Stryker, 2012*; *Magueresse and Monyer, 2013***), but whether and how it shapes the over-representation of cardinal orientation is not known. In this section, we study the changes in tuning properties of inhibitory PV+ interneurons during the visual development.

In the network model, we observe structural changes in the tuning properties of inhibitory neurons during learning. First, the inhibitory neurons show a modest level of orientation tuning at the beginning, which is lost during learning (***Figure 4***A-B). On the population level, the orientation selectivity index significantly decreases during learning (***Figure 4***C, (two-sample Kolmogorov–Smirnov test, p < 2.2E-16)). Since PV+ inhibitory neurons receive dense and unselective connection from the excitatory population, changes in their orientation selectivity may reflect changes in the distribution of preferred orientations as well as population tuning curves of excitatory neurons (***Runyan and Sur, 2013***).

**Figure 4.**
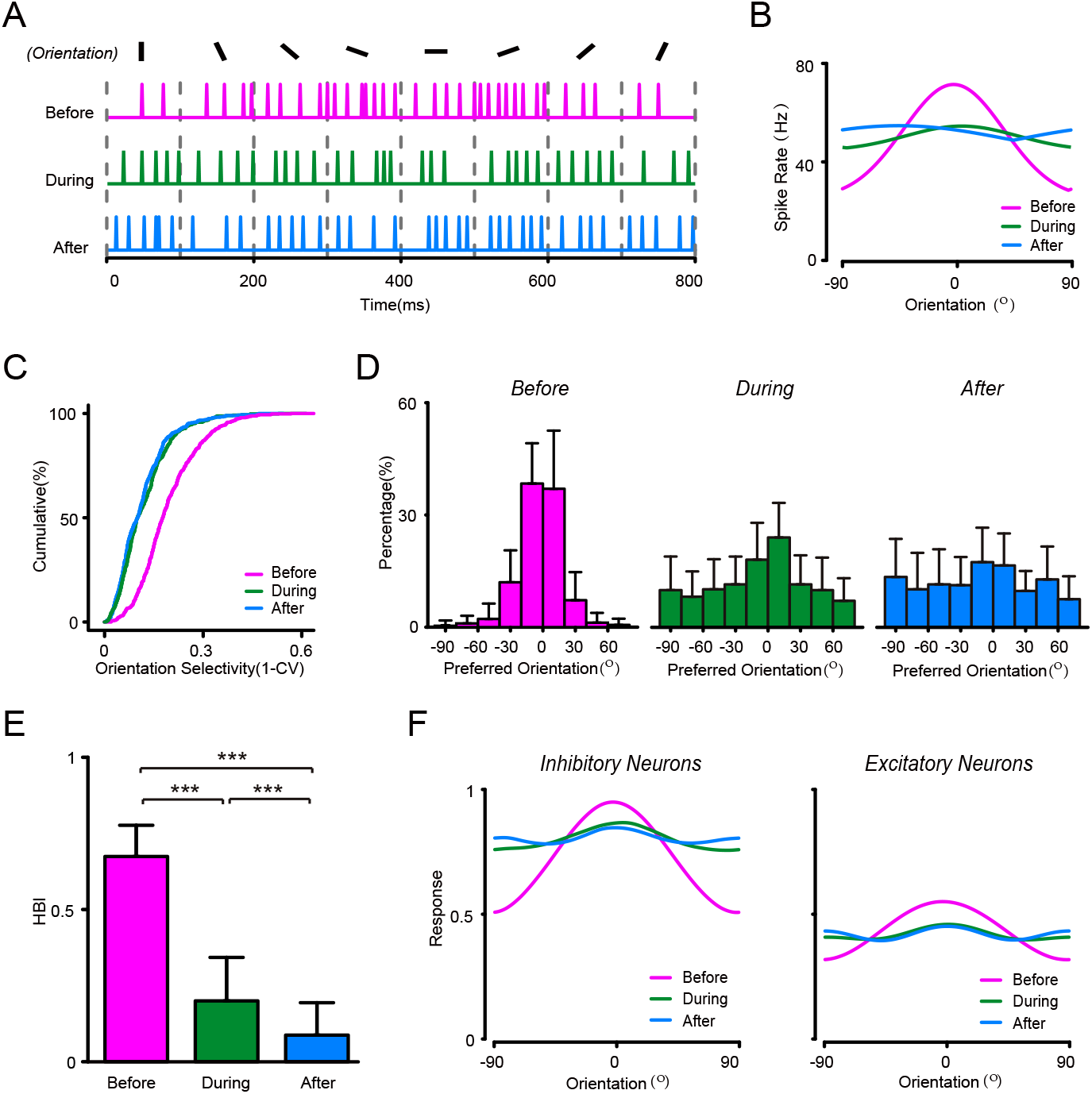
Simulated inhibitory neurons provide competitive inhibition during learning **A**. Spike trains of a representative simulated inhibitory neuron to stimuli with 8 different orientations, at different learning stages, before (magenta), during (green) and after (blue) learning. **B**. Orientation tuning curves of a representative inhibitory neuron at different learning stages (the same neuron as in **A**). **C**. Distribution of global orientation selective index (gOSI) of simulated inhibitory neurons at different learning stages. **D**. Distribution of preferred orientation of simulated inhibitory neurons before (left), during (middle), and after (right) learning. **E**. HBI of simulated inhibitory neurons at different learning stages. **F**. Population tuning curve of simulated excitatory (left) and inhibitory (right) populations at different learning stages. P values in **E**. are calculated by the Kolmogorov–Smirnov test. *** indicates P<0.01; ** indicates 0.01<P<0.05.

Second, inhibitory neurons provide non-uniform inhibition during the early learning stage, most prominently at horizontal orientations. The distribution of preferred orientation of the inhibitory population shows a horizontal bias at beginning of the learning period, with more neurons preferring horizontal orientation (***Figure 4***D left), which are also confirmed by HBI (t-test between before and during learning, p < 2.2E-16; t-test between before and after learning, p < 2.2E-16, ***Figure 4***E). Moreover, the activity of inhibitory population, represented by the population tuning curve, shows higher activity at horizontal orientation, matching the population activity of excitatory neurons (***Figure 4***F magenta lines). As learning proceeds, the population tuning curve as well as the distribution of preferred orientation of inhibitory neurons become more isotropic, as the horizontal bias disappears in the excitatory population. We propose that non-uniform recurrent inhibition provided by inhibitory neurons plays a key role in the emergence of probabilistic representation in the network.

Our model suggests that inhibitory neurons provide non-uniform inhibition at early learning stage, which becomes more isotropic at later stage. Here, we examine whether such phenomenon exists in PV+ inhibitory interneurons, the major players of feedback inhibition in cortical circuits (***Hu et al., 2014***). We implement *in vivo* two-photon laser imaging guided cell-attached recording to measure spiking response specifically from the PV+ interneuron in the V1 binocular area, responding to the same drifting gratings with 6 different orientations (***Figure 5***A). To examine the developmental changes, we record PV+ interneurons in the mice shortly after the EO (P17-18), during the CP (P27-28) and in the adult stage (P55-56), respectively (***Figure 5***B). By calculating the OSI, we find that the orientation tuning of PV+ interneurons is stronger at the period shortly after the OE than that of the other two later stages (***Figure 5***C). The OSI amplitudes of PV+ interneurons are decreased significantly from the OE to the CP (two-sample Kolmogorov–Smirnov test between P17-18 and P27-28, p = 1.47E-7), and they continuously remain low through the adult stage (***Figure 5***D, two-sample Kolmogorov– Smirnov test between P17-P18 and P55-56: p = 2.93E-8). These results largely agree with previous experimental findings that the inhibitory PV+ interneurons exhibit an initially strong orientation tuning shortly after the OE (***Kuhlman et al., 2011*; *Li et al., 2012***).

**Figure 5.**
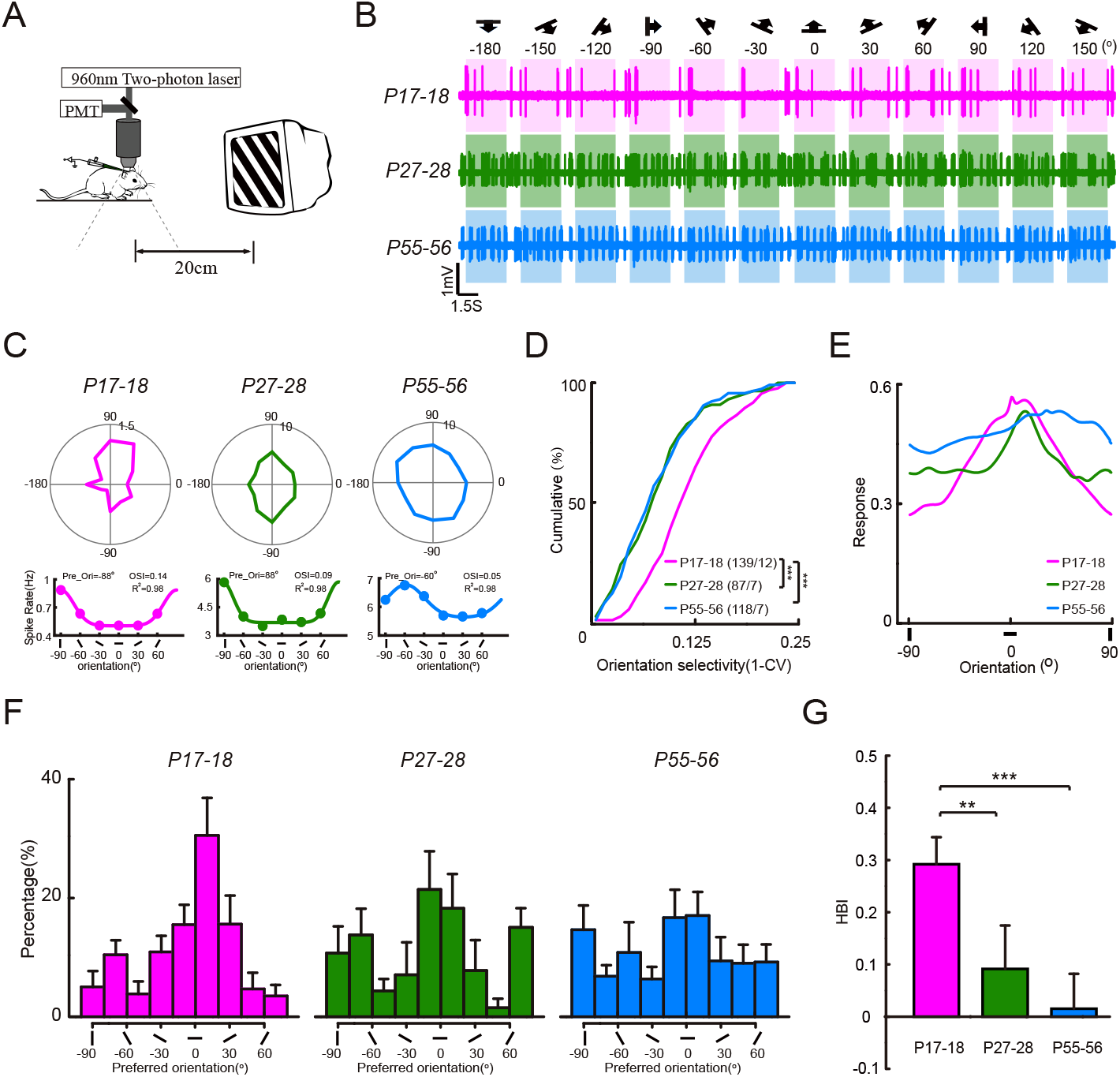
The orientation selectivity and the preferred orientation bias of PV neurons in mice V1 are likely to change at different postnatal stages. **A**. Schematic diagram illustrating *in vivo* two-photon imaging guided cell-attached recording with glass micropipettes (green, Alexa 488) of spike responses from layer 2/3 PV neurons in mouse V1. **B**. The responses of representative neurons at different postnatal stages to presented drifting gratings with 12 different directions (magenta trail, P17-18; green trail, P27-28; blue trail, P55-56). The stimulus directions and orientations are indicated by the symbols shown on top. **C**. Top, the orientation-tuned responses of representative neurons in **A**. Bottom, the curves denote the method to use one-peak Gaussian function fitting to detect the preferred-orientation of representative neurons in **A. D**. Cumulative distributions of gOSI (1-CV) values of PV+ neurons at 3 different postnatal stages (magenta line, P17-18; green line, P27-28; blue line, P55-56). **E**. Population tuning curves of PV+ neurons at 3 different postnatal stages (magenta line, P17-18; green line, P27-28; blue line, P55-56). **F**. Distributions of preferred orientations of PV+ neurons in P17-18 mice (126 cells / 12 mice), P27-28 mice (74 cells / 7 mice) and P55-56 mice (100 cells / 7 mice). **G**. HBI of PV+ neurons in 3 different postnatal stages. The P values in **D** and **G** are calculated by the Kolmogorov–Smirnov test and t-test respectively. *** indicates P<0.01; ** indicates 0.01<P<0.05.

We further analyze the distribution of preferred orientation of PV+ interneuron at different developmental stages. Strikingly, we find that their preferred orientation distributions also shows an apparent bias to the horizontal orientation at the earlier stage and such bias gradually disappears in the later development (***Figure 5***F), a process highly assemble to the observations in the PC population (***Figure 3***). The horizontal bias measured HBI is highest for PV+ interneuron shortly after EO, and decreases during the CP (t-test, p = 0.032) and adult stage (t-test, p = 0.001, ***Figure 5***G). Consistently, the population tuning curves of PV+ interneurons show a horizontal bias shortly after the EO, and then become absent in the later development ***Figure 5***E). Taken together, these experiment results strongly suggest that PV+ interneurons can provide non-uniform inhibition in the cortical network during the early cortical development, which endorse the notion raised by the model simulations.

### Non-uniform feedback inhibition is necessary for the emergence of the probabilistic representation

Our model and experiment results suggest that inhibitory neurons provide non-uniform feedback inhibition during development, but how it contributes to the visual development is not known. We hypothesize that the non-uniform feedback inhibition is necessary for the emergence of probabilistic representation in the cortical network for following reasons: first, different from WTA inhibition, feedback inhibition provides a softer form of inhibition in the cortical network (***Avermann et al., 2012*; *Hofer et al., 2011***), where the level of inhibition is proportional to the level of excitation (***Isaacson and Scanziani, 2011***). This softer form of inhibition allows a subset of excitatory neurons to respond to a specific input feature, which is ideal for representing a probability distribution in the network. Second, when applied with the STDP learning rule, the feedback inhibition induces excitatory neurons to compete for the right to respond to input patterns, enabling a powerful learning paradigm referred as “competitive learning” (***Grossberg, 1976*; *Masquelier et al., 2009*; *Nessler et al., 2013***). As a result, the distribution of preferred stimuli has an appropriate shape respective to the input prior probabilities: if prior probability of an input pattern is higher, there will be a larger fraction of neuron preferring that pattern; however, if an input pattern is overrepresented in the network, the level of inhibition will rise and prevent excitatory neurons from learning that pattern. In other words, the network can be described by an adaptive k-WTA computation (***Lange, 1992***), where k is adaptively determined by the prior probabilities and the level of inhibition in the network.

To verify our hypothesis, we change the non-uniform feedback inhibition in the network to uniform inhibition, by ablating the recurrent connections from excitatory neurons to inhibitory neurons (***Figure 6***A). We also increase the feed-forward connection weight received by inhibitory population to keep their mean firing rates unchanged. As a result, the uniform feed-forward activity in the network is independent from the activity of excitatory population (***Figure 6***B). Under such condition, the excitatory population fails to transform from horizontal bias to cardinal bias (***Figure 6***C). Our results suggest that the non-uniform feedback inhibition, rather than uniform feed-forward inhibition, is necessary for the emergence of probabilistic representation in the cortical network.

**Figure 6.**
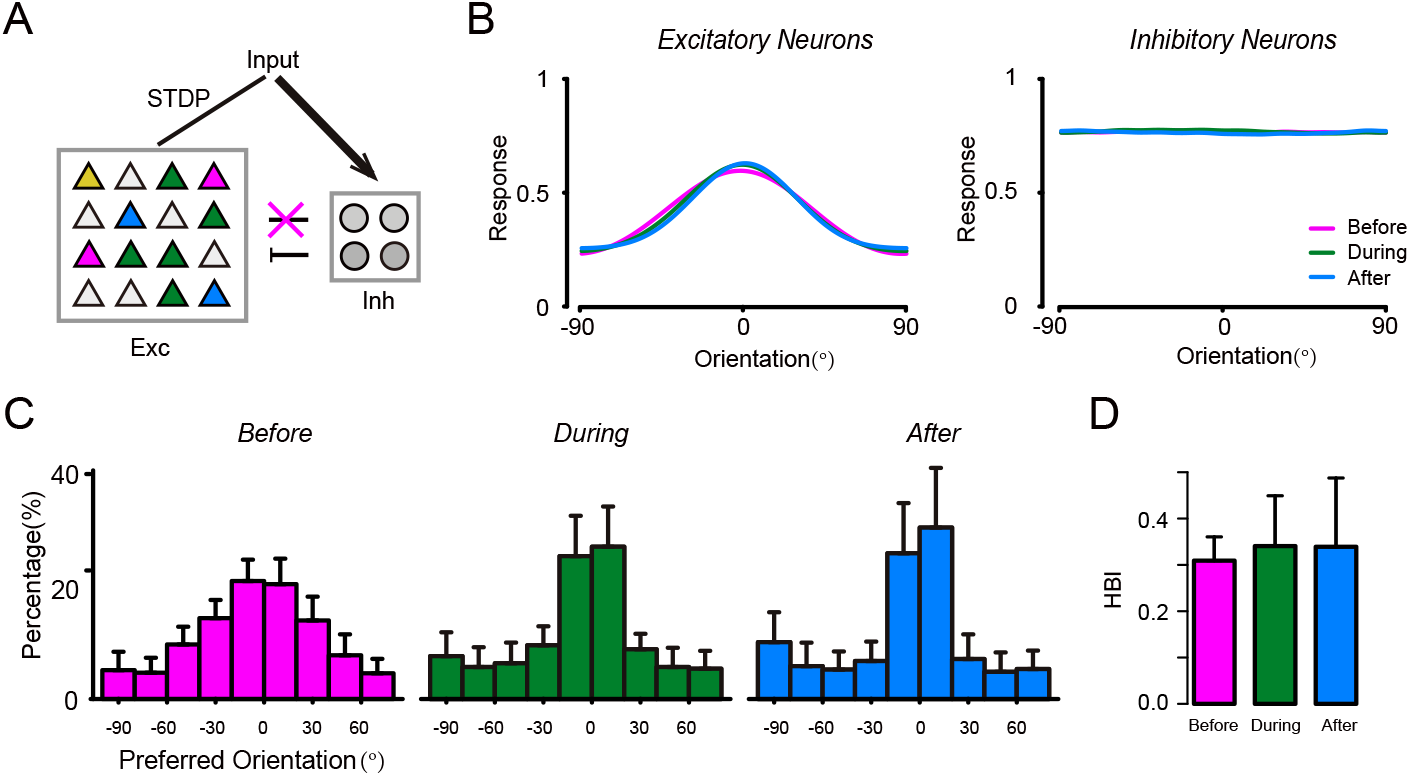
Failed emergence of probabilistic representation in the absence of competitive inhibition. **A**. Schematic diagram of the network structure with ablation of connections from cortical excitatory neurons to inhibitory neurons. **B** Population tuning curve of excitatory (left) and inhibitory (right) populations at different learning stages. magenta, green and blue lines indicate before, during and after learning. **C**. Distribution of preferred orientation of excitatory neurons before (left), during (middle), and after (right) learning. **D**. HBI of simulated excitatory neurons at different learning stages.

### Analysis using a simplified rate model

Our theoretical and experimental results suggest that the prior probability and non-uniform inhibition are essential to generate the probabilistic representation in the cortical network. To further demonstrate this point, we construct a simplified rate model to elucidate the effects of the prior probability and non-uniform inhibition on the dynamics of synaptic plasticity.

In the simplified model, we consider a network of N mutually inhibiting neurons with linear dynamics, and they receive input patterns **x** with synaptic weights **w**_**i**_ = {*w*_1*i*_, *w*_2*i*_} (***Figure 6***A). The feed-forward synaptic weights are modified according to the BCM learning rule (***Bienenstock et al., 1982***), in which the activities of post-synaptic neuron have a non-linear effect on the changes of synaptic weights (***Figure 7***B). In our case, we consider 2 input patterns with different priors, i.e., ***P*** (*x*^(1)^) = *ρ* and ***P*** (*x*^(2)^) = 1 − *ρ*. The effect of non-uniform inhibition is captured by the ratio of the inhibition level under different input patterns (***L***^(1)^/***L***^(2)^), which is influenced by the proportions of neurons preferring different patterns. Previous studies have analyzed how the prior probability shapes the attractor properties in a similar network (***Cooper and Scofield, 1988*; *Intrator and Cooper, 1992*; *Udeigwe et al., 2017***), but phase plane analysis has not been applied and the effect of non-uniform inhibition has not been explored.

**Figure 7.**
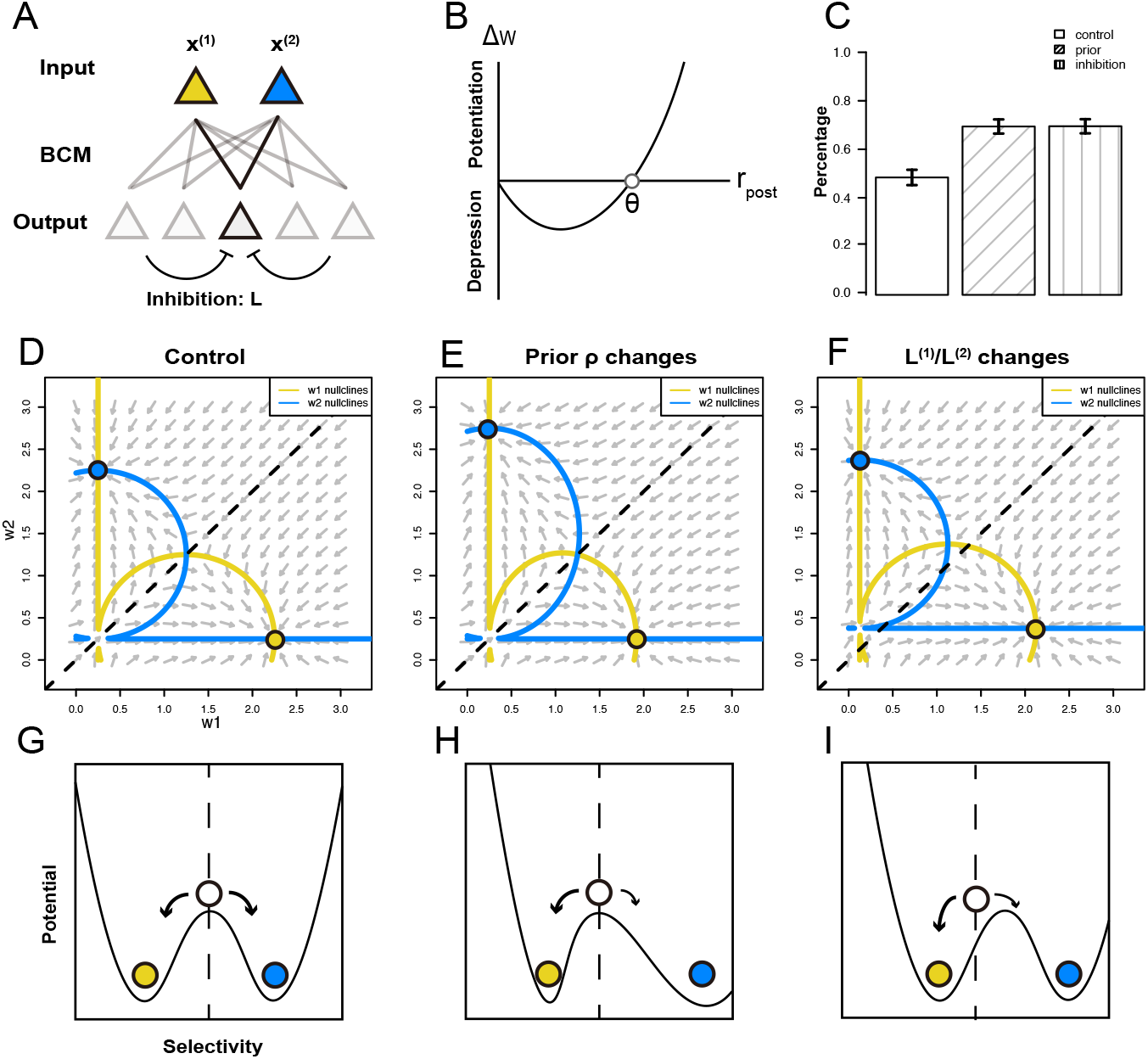
Analysis using a simplified rate model. **A**. Schematic diagram of the simplified rate model. **B**. BCM learning rule used in the simplified model. **C**. Effects of the prior and non-uniform inhibition on the synaptic weights of a non-selective neuron. **D-F**. Phase plane of the feed-forward synaptic weight for control group (**D**), when the prior probability of pattern 1 increases (**E**) or the inhibition ratio decreases (**F**). **G-I**. Schematic diagrams of the potentials governing the dynamics of the synaptic weights for groups in **D**-**F**. Yellow and blue circles represent attractor states that are selective to pattern 1 and 2 respectively, and white circle represents the non-selective state.

We focus on inspecting the synaptic weights of a non-selective neuron, while keep the weights of other neurons fixed (by setting their weights near to the attractors). This implies that the ratio of inhibition levels under different input patterns (***L***^(1)^/***L***^(2)^) are held constant during learning. First, we study the dynamics of synaptic weights under three different conditions: (1) uniform prior and uniform inhibition: *ρ* = 0.5 and ***L***^(1)^/***L***^(2)^ = 1, (2) non-uniform prior and uniform inhibition *ρ* = 0.6 and ***L***^(1)^/***L***^(2)^ = 1, (3) uniform prior and non-uniform inhibition: *ρ* = 0.5 and ***L***^(1)^/***L***^(2)^ = 1/2. Under uniform prior and uniform inhibition, the neuron has equal chances to become selective to either pattern (***Figure 7***C). But under non-uniform prior or non-uniform inhibition, the chances are no longer equal: the neuron has a higher chance to become selective to the pattern with the higher prior or the lower level of inhibition (***Figure 7***B). To further unveil the effects of non-uniform prior and feedback inhibition on the network dynamics, we perform phase plane analysis on the synaptic weights from different input units. For uniform prior and uniform inhibition, the phase plane is symmetrical to the diagonal line (***Figure 7***D), indicating that the neuron has equal chances to become selective to either pattern. But when the prior probability becomes non-uniform, e.g., *ρ* increases, the attractor that is selective to pattern *x*^(1)^ becomes closer to the diagonal line, while the attractor to *x*^(2)^ moves away (***Figure 7***E). In such a case, the neuron has a higher chance to become selective to *x*^(1)^. When the feedback inhibition becomes non-uniform, e.g., ***L***^(1)^/***L***^(2)^ decreases, the basin boundary separating the two basins of attraction shifts towards upper-left, which has the similar effect as increasing the prior probability (***Figure 7***F). In summary, our analysis shows that non-uniform prior and feedback inhibition shape the changes of synaptic weights, which affect the selectivities of neurons in the network (***Figure 7***G-I).

### The learned internal representation supports Bayesian-like inference

Learning and inference are not separable in the brain (***Friston, 2003*; *Fiser et al., 2010***). However, the validation method of previous biological plausible learning model solely concentrated on comparing the “receptive field” properties of model units with those of neurons. And how the learned model contributes to inference is left implicit (for exceptions, please see ***Rao and Ballard*** (***1999***); ***Nessler et al***. (***2013***)). Here, we show that our network is able to perform Bayesian-like inference based on the learned internal representation, where the posterior of the orientation can be read out by a population vector decoder (***Georgopoulos et al., 1986***).

To decode the orientation from neural activity, we construct a encoder-decoder model based on the network described in the previous section (***Girshick et al., 2011*; *Fischer and Peña, 2011***). The network encodes the ground truth orientation *θ* in the population activity, from whom an estimate of the orientation 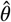 is decoded by a population vector decoder (***Figure 6***A). The population vector decoder computes the sum of the directional vectors associated with the preferred orientation for each neuron, weighted by its firing rates (***Figure 8***B). Previous studies show that if the distribution of preferred stimuli matches the prior probabilities of the stimuli, and the tuning curves of neurons follow Gaussian shapes, the population vector will be consistent with the Bayesian estimate of the stimulus (***Fischer and Peña, 2011*; *Girshick et al., 2011*; *Ganguli and Simoncelli, 2014***). Because the distribution of preferred orientation shift from horizontal bias to cardinal bias during development (***Figure 8***C), we expect the decoded orientations to have different Bayesian characteristics in terms of estimation variability and bias across different development stages.

**Figure 8.**
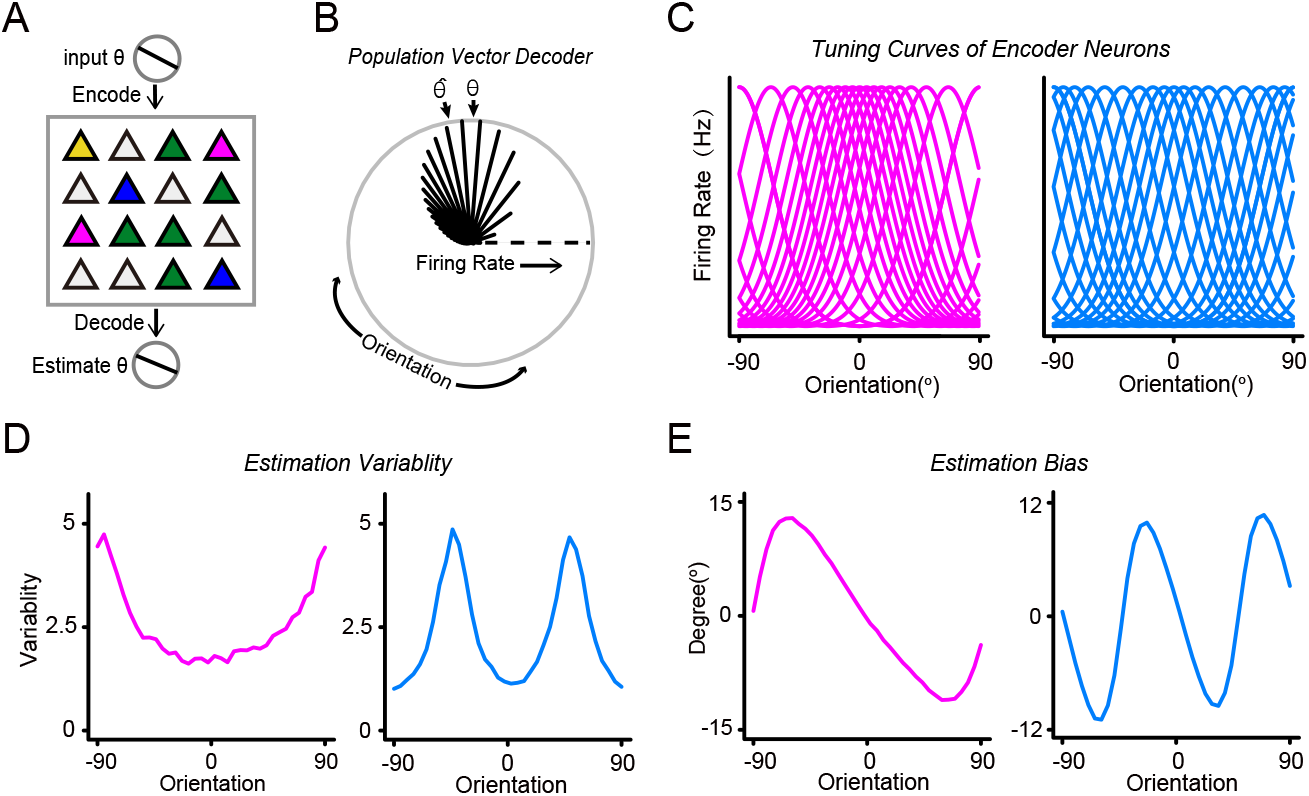
Bayesian-like inference from the probabilistic representation of the network. **A**. Schematic diagram of the neural encoder-decoder model. Orientation input is encoded by the neural network with neural response. Then the orientation is estimated by the population vector decoder. **B**. Schematic diagram of the population vector decoder with a non-uniform neural population. **C**. Tuning curves of the encoder neural population with non-uniform orientation preferences before (left) and after learning (right), the same neural population as in Fig.2 (only a subset of neurons shown). **D**. Variability of the estimated orientations before (left) and after learning (right). **E**. Bias of the estimated orientations before (left) and after learning (right)

The variability of the decoded orientation is computed as the *s*.*d*. of the estimated orientation given the ground-truth orientation, which is relevant to the discrimination thresholds in psychophysics (***Green et al., 1966***). Before learning, the variability is lowest at horizontal orientation and highest at vertical orientation, which suggests that the discriminability is best at horizontal orientation and worst at vertical orientations for young mice (***Figure 8***D, left). While after learning, the variability is lowest at cardinal orientations and highest at oblique orientations. Thus, the results suggest that the discriminability of adult mice is best at cardinal orientation and worst at oblique orientations (***Figure 8***D, right), which is consistent with the “oblique effect” observed in monkeys and human performing orientation discrimination tasks (***Appelle, 1972*; *Bauer et al., 1979***). In general, the discriminability is the best at the peak of the distribution of preferred orientation of the encoding population. Recent researches show that mice are able to discriminate horizontal and vertical orientations (***Long et al., 2015*; *You and Mysore, 2020***), but their discriminant thresholds for different orientations have not been tested systematically.

The non-uniform distribution of preferred orientation in the encoding population will cause bias in estimation (***Girshick et al., 2011*; *Wei and Stocker, 2015***). The perceptual bias is computed as the expected difference between the estimated and ground-truth orientations, conditioned on the ground-truth orientation. In our model, the bias shows a “prior attraction” effect, where the prior distribution leads to an attraction shift of the estimator (***Figure 8***B) (***Girshick et al., 2011***). For the network before learning, the perceptual bias is unimodal, indicating that orientations are perceived to be oriented closer to the horizontal orientation (***Figure 8***E, left). And the bias is 0 at horizontal and vertical cardinal orientation, and as large as 12° at oblique orientations. For the network after learning, the perceptual bias is bimodal, indicating that orientations are perceived to be oriented closer to the nearest cardinal orientation (***Figure 8***E, right). The bias is 0 at cardinal and oblique orientations, and as large as 10° in between. Recent experiment shows neuronal representation in V1 of adult mice affects their behavior bias (***Jin and Glickfeld, 2019***). In summary, the cortical network supports Bayesian inference based on the learned representation.

## Discussion

In this work, we combine mathematical modeling with electrophysiology experiment to study the emergence of probabilistic representation in cortical network of mice V1 during development. We find that PV+ inhibitory neurons played an important role in the emergence of probabilistic representation in V1 through feedback inhibition. With electrophysiology recording, we find a novel and tight relationship between the orientation tuning of PCs and PV+ inhibitory neurons across different development stages. Furthermore, ablation study of the network model shows that the non-uniform feedback inhibition is necessary for the emergence of probabilistic representation. Lastly, we show that the network supports Bayesian-like inference based on the learned internal representation. In summary, our results support the proposal that the microcircuit consisted of PCs and PVs inhibitory interneurons might be the building block for Bayesian inference (***Darlington et al., 2018***).

The orientation tuning properties of the PV+ inhibitory interneurons change structurally during the visual development, which may be crucial for sharpening the orientation selectivity and initiating the CP of excitatory binocular plasticity in the developing visual cortex (***Kuhlman et al., 2011*; *Li et al., 2012***). PV+ inhibitory interneurons are orientation selective at early development stages (few-days after EO) and become unselective before onset of CP. Since the orientation selectivity of PV+ inhibitory neuron tends to reflect net biases of the surrounding neurons (***Kerlin et al., 2010*; *Packer and Yuste, 2011***), we propose such structural change in PV+ inhibitory interneurons reflects the shift in preferred orientation of the PCs during development: few days after EO, the local PC population is biased towards horizontal orientations, thus the PV neurons have high orientation selectivity and are likely to prefer horizontal orientations (***Figure 6***A); while at later stage of development, the horizontal bias in PC population is lost, and the number of PCs preferring to a specific orientation and its orthogonal orientations are likely to be equal, resulting a lower orientation selectivity in PV+ inhibitory neurons (***Figure 9***B).

Primary sensory cortex may simultaneously employ different strategies to represent the uncertainty. Besides the framework of probabilistic representation that we currently studied, where the prior distribution is embedded in the distribution of preferred stimuli (***Fischer and Peña, 2011*; *Girshick et al., 2011*; *Ganguli and Simoncelli, 2014***), theorists have proposed other frameworks for probabilistic representation in the primary sensory cortex. ***Berkes et al***. (***2011***) proposed that the spontaneous activity of PCs can represent the prior distribution of the stimuli, while the stimulus-evoked activity can represent the posterior. Their framework is supported by experiment data in ferrets V1, where the spontaneous activities reflect prior expectations of the evoked activities by natural scenes. Probabilistic population code (***Ma et al., 2006***) proposes that the uncertainty of the stimulus can be encoded by the gain of the population tuning curve. Recent experiments (***Walker et al., 2020***) suggest that tuning widths of the likelihood function decoded from V1 explain the behavior variability, which partly support the probabilistic population code. But the circuit mechanism for acquiring above probabilistic representations are not understood and should be investigated in future research.

Our current research focus on the developmental role of the major player of feedback inhibition, the PV+ inhibitory neurons, but other types of inhibitory neurons may also regulate the emergence of probabilistic representation in V1. Similar to PV+ neurons, somatostatin (SOM) inhibitory neurons have dense and reciprocal synaptic connections with near-by PCs (***Fino and Yuste, 2011***). However, compared to PV neurons, the inhibition provided by SOM neurons is more facilitatory (***Gentet et al., 2012*; *Miao et al., 2016***), more orientation selective (***Ma et al., 2010*; *Tremblay et al., 2016***), and targeting to distal dendrites of PCs. Thus, SOM neurons may regulate the visual development through a more accurate form of inhibition that gates the feed-forward inputs received by PCs. Moreover, VIP inhibitory neurons regulate visual development through dis-inhibition. A recent study showed that developmental dysfunction of VIP interneurons would eliminate PCs’ bias toward horizontal visual stimuli (***Batista-Brito et al., 2017***). The emergence of probabilistic representation may involve the inhibitory circuits consisting various subtypes of inhibitory neurons.

We provide several testable predictions: First, we predict that the feedback inhibition is necessary for the emergence of overrepresentation of cardinal orientations during normal visual development. This prediction can be tested by lowering the activity of PV+ inhibitory interneurons, possibly by knocking down PV or GAD65 gene expression in V1. Second, we predict that, as a consequence of non-uniform distribution of preferred orientation in neural population, the discrimination threshold of mice is non-uniform: lowest at horizontal orientations for mice few-days after EO, and lowest at cardinal orientations for adult mice. This can be directly tested on mice with orientation discrimination tasks (***Long et al., 2015*; *You and Mysore, 2020***).

Learning, representation and inference are not separable in the brain. Our work suggests that the feedback inhibition circuit shapes the emergence of probabilistic representation and supports Bayesian-like inference, which may serve as a building block for Bayesian inference in the brain.

## Methods and Materials

### Spiking Neural Network of Mouse V1

The circuit model consists of a two-layer neural network. The first layer is the input layer with *N*_*P*_ Poisson neurons. The second layer represents the cortical network, which contains *N*_*E*_ excitatory and *N*_*I*_ inhibitory neurons, mimicking the interacting populations of pyramidal cells and PV+ inhibitory neurons in mice V1.

#### Cortical neurons

The cortical layer is modeled by a recurrent network of *N* = 100 leaky integrate-and-fire neurons, of which 80% are excitatory and 20% are inhibitory. The sub-threshold dynamics of membrane potential *u*_*i*_ of neuron *i* follows

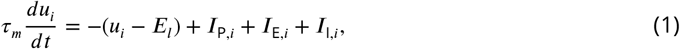

where the membrane time constant *τ*_*m*_ = 20 ms and the resting potential *E*_*l*_ = −70 mV. *I*_P,*i*_ describes the current received from the input layer, *I*_E,*i*_ and *I*_I,*i*_ are the recurrent inputs from cortical excitatory and inhibitory neurons. When the membrane potential reaches the threshold *u*_*th*_ = −50 mV, a spike is generated and transmitted to all post-synaptic neurons, then the membrane potential is reset to the resting potential *u*_0_ = −70 mV.

#### Input

The cells in the input layer are modeled as Poisson neurons with time-varying firing rates, which depends on the 1-dimensional stimulus. The continuous orientation pattern *θ* ∈ [0, *π*] is encoded by *N*_*P*_ = 100 Poisson neurons using a positional code with a Gaussian profile, illustrated in ***Figure 6***A.

The firing rate of input neuron *i* in response to a stimulus at orientation *θ* at time t is:

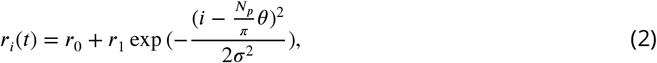

where *r*_0_ and *r*_1_ is spontaneous and maximum firing rate of the input neuron, *σ* is the spreads of the Gaussian function (***Song and Abbott, 2001***). Periodic boundary conditions are imposed to avoid edge effects.

The prior probabilities of the input patterns are described in the simulation detail sections.

#### Network connectivity

The excitatory neurons are reciprocally and randomly connected with inhibitory neurons, with connection probabilities *p*_EI_ = *p*_IE_ = 50%. Inbitory neurons are recurrently connected with each other with the probability *p*_II_ = 75%. The recurrent connections between excitatory neurons are not included in the model for simplicity, but simulations with the recurrent connections included do not change our results. The connection probability are based on the connectivity data of pyramidal cells with PV^+^ inhibitory neurons in V1 from (***Miao et al., 2016***). Both excitatotry and inhibitory neurons receive feed-forward connections from the input layer with probabilities *p*_PE_ = 100% and *p*_PI_ = 40%.

#### Synaptic currents

The synapses from input neurons to cortical neurons are all excitatory, with reversal potential *V*_*E*_ = 0 mV, synaptic constant *τ*_syn_ = 3 ms, and synaptic conductance *g*_PE_ and *g*_PI_ for excitatory and inhibitory populations. The input-cortical synaptic current *I*_P,*i*_ received by cortical neuron *i* in neural population A ∈ {E, I} is

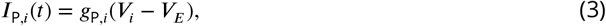

With 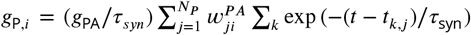, where 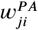 is the weight of the input-cortical connection from input neuron *j* to cortical neuron *i*, and *t*_*k,j*_ is the time of the *k*-th spike generated by input neuron *j*.

The cortical excitatory and inhibitory synaptic currents have similar forms as that of the input synaptic current. The cortical excitatory input received by neuron *i* in population A ∈ {E, I} is

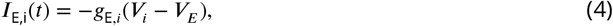

With 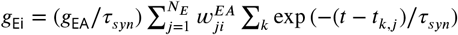, and the synaptic conductance *g*_EI_ = 0.06.

The inhibitory input received by neuron *i* in population A ∈ {E, I} is

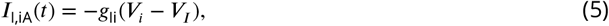

With 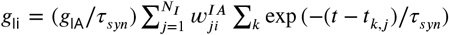, where synaptic conductance *g*_IE_ = 0.02 and *g*_II_ = 0.02

#### Plasticity rule

The connection weights from input neurons to excitatory neurons are plastic and modified by the standard STDP rule illustrated in ***Figure 1***B (***Song and Abbott, 2001***). The connection weights *w*_*ij*_ are updated as follows. For each postsynaptic spike at time *t*_post_, only nearest-neighbor preceding presynaptic spike is considered. For each such pre-before-post spike pair with time difference *t*_post_ − *t*_pre_, the weight is potentiated by

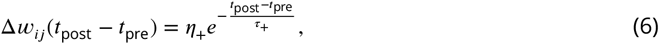

with time constant *τ*_+_ = 14 ms and learning rate *η*_+_ = 0.005. For each presynaptic spike at time *t*_pre_, only nearest-neighbor preceding postsynaptic spike is considered. For each such post-before-pre pair with time difference *t*_pre_ − *t*_post_, the weight is depressed by

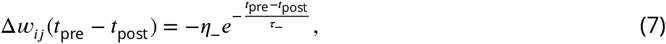

with time constant *τ*_+_ = 34 ms and learning rate *η*_−_ = 0.0022 (keeping 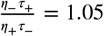).

To avoid unbounded weight growth, we introduce a fast homeostatic plasticity. We keep the sum of the feed-forward weights received by an excitatory neuron 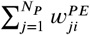 to be constant at its initial value, by rescaling weights after each update. Negative or unconnected synaptic weights are set to be zero. Weights of non-plastic connections, which keep constant during simulation, are set to be 1 if two neurons are connected, and 0 otherwise.

#### Population vector decoder

We use a population vector decoder to decode the orientation from the network response. The population vector is computed as a linear combination of the preferred orientation vectors of the neurons, weighted by the firing rates given the input orientation *θ*

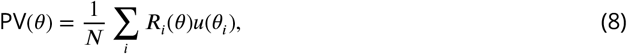

where *u*(*θ*_*i*_) is a unit vector pointing in the *i*^th^ neuron’s preferred orientation and _*i*_(*θ*) is the firing rate of the *i*^th^ neuron, drawn either from a Poisson or Gaussian distribution as described above. The estimated orientation 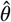 is found by computing the direction of the population vector using the inverse tangent.

### Simplified Rate Model

In our simplified rate network model, we consider a network of *N* mutually inhibiting neurons and input units representing patterns (***Cooper and Scofield, 1988*; *Intrator and Cooper, 1992***). The activities of neurons follow linear dynamics:

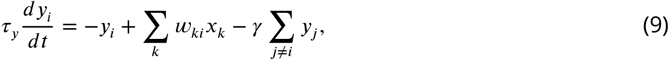

where _*i*_ is the firing rate of neuron *i*, _*k*_ is the activity of input unit *k, w*_*ki*_ is the synaptic weight from input unit *k* to neuron *i*, is the lateral inhibition parameter, and *τ*_y_ is the time constant. The synaptic weights evolves according to the BCM rule as (***Bienenstock et al., 1982***):

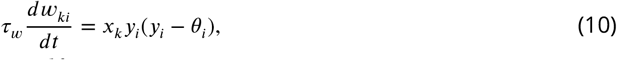

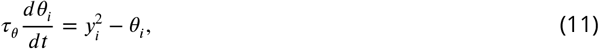

where *θ* is the “sliding threshold” that models homeostatic process, and *τ*_*w*_ and *τ*_*θ*_ are time constants. The system can be further simplified by assuming that the firing rate and homeostatic processes have faster dynamics than the synaptic plasticity (*τ*_*y*_, *τ*_*θ*_ ≪ *τ*_*w*_), which gives

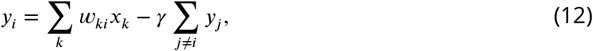

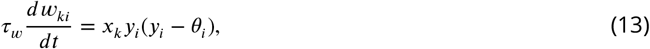

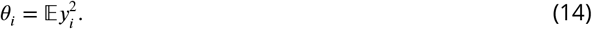

For illustration purpose, we choose two orthogonal input patterns (= 2), with ^(1)^ = [1, 0] and ^(2)^ = [0, 1] (for correlated input patterns, please see ***Udeigwe et al***. (***2017***)). The probabilities of presenting ^(1)^ and ^(2)^ are and 1 − respectively. This learning system is stochastic in nature. To gain more insight, we can transform the stochastic dynamic system to a deterministic one using the mean-field approach, by studying the average effect of the probabilistic input on the synaptic weight (***Cooper and Scofield, 1988*; *Intrator and Cooper, 1992*; *Udeigwe et al., 2017***). A mean field equation for the synaptic weights is:

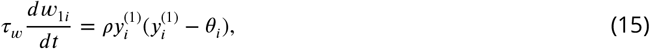

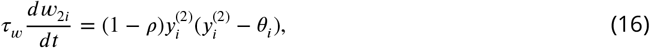

where 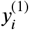 and 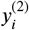 are the firing rates of neuron *i* under input pattern 1 and 2 respectively, and 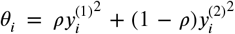. Similarly, the mean field equation for the firing rates of neuron *i* as a consequence of synaptic plasticity is:

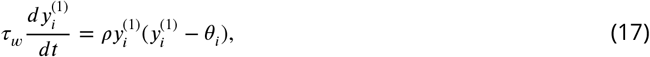

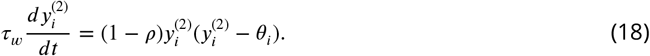

Above equations allow us to analyze the effect of prior probabilities and non-uniform feedback inhibition using phase plane analysis. By setting above derivatives to 0, we get the fixed points of the dynamic system: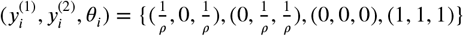. The fixed points 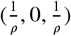 and 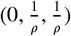 are stable, while (0, 0, 0) and (1, 1, 1) are neither stable nor selective (***Castellani et al., 1999***). We can study the effects of prior probabilities and non-uniform feedback inhibition on the dynamics of synaptic weight of neuron *i* by changing *ρ* and 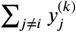 respectively.

### Details to computer simulations of the network model

In this section, we provide details to the computer simulations reported in Results. All simulations were performed in R, with a discretization time step Δ*t* of 1 ms. Each simulation is repeated for 25 times. All codes are available at https://github.com/AmazingAng/EmergeProbV1.

#### Details to simulations for *Figure 1*

For ***Figure 1***C-D, the network receives continuous orientation inputs, mimicking the effect of strips rearing, where the mice have restricted visual experience with only one orientation (***Kreile et al., 2011***). The prior probability of orientation inputs follows a von-Mises distribution with center at the experienced orientation (-90°, -45°, 0°, 45°) and concentration = 0.5 (see ***Figure 1***C). Before training, the feed-forward connection weights of excitatory neurons are set according to a Gaussian profile:

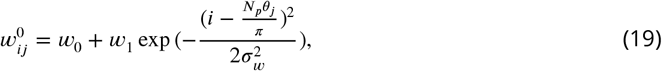

where *w*_0_ and *w*_1_ are the minimum and gain factor of the initial weight, *σ*_*w*_ is the spread of the Gaussian function, *θ*_*j*_ is the initial preferred orientation of neuron *j* (***Song and Abbott, 2001***). To mimicking the over-representation of cardinal orientations in mice V1, the initial preferred orientation *θ*_*j*_ is uniformly and randomly distributed in the range of 0 to *π*. During training, each pattern is selected randomly according to its prior probability, corrupted with white noises with zero mean and 15° s.d., and presented for 100 ms. The training period lasts for 1500 s. The tuning curve properties of excitatory neurons in the network are evaluated before and after learning. The synaptic conductance between inputs to cortical neurons are *g*_PE_ = 0.012 and *g*_PI_ = 0.006

#### Details to simulations for ***Figure 2*** and ***Figure 4***

We examine the emergent properties of the cortical network model on an input distribution that mimics the distribution of orientations in natural images. Before training, the distribution of preferred orientations *θ*_*j*_ is initialized following a von-Mises distribution with center = *π*/2 and concentration = 0.8, according to the over-representation in horizontal orientations in mice V1 after EO (***Hagihara et al., 2015***).

During the training phase, the feed-forward connection weights of excitatory neurons are modified by the STDP learning rule. Orientation inputs are drawn randomly according to the local orientation distribution measured in photography *p*(*θ*) = (2 − |sin(2*θ*)|) illustrated in ***Figure 6***A (***Girshick et al., 2011*; *Wei and Stocker, 2015***). To measure the orientation tuning of neurons, during the testing phase, STDP in the network is disabled, and the network is tested with 8 orientation steps, each for 1000 ms (see ***Figure 2***B and ***Figure 6***A). Other conditions are the same as ***Figure 1***.

#### Details to simulations for ***Figure 6***

We examine whether competitive inhibition provided by inhibitory neurons is necessary for the emergence of probabilistic representation in the network. For this purpose, we ablate the synaptic connection from cortical excitatory population to inhibitory population, and increase the synaptic conductance from input neuron to inhibitory neuron *g*_PI_ from 0.006 to 0.03 (see ***Figure 6***A). This will remove all tuned inputs received by inhibitory neurons, while keeping their mean firing rate intact. Other settings are the same as ***Figure 2***.

#### Details to simulations for ***Figure 8***

***Figure 8*** shows that the network performs Bayesian-like inference with population vector decoder, on the embedded probabilistic representation. Theoretical studies show that if the probability distribution of preferred stimuli matches the prior probability of the stimuli, and the tuning curves are proportional to the likelihood function, the population vector will be consistent with the Bayesian estimate of the stimulus (***Fischer and Peña, 2011***).

The network is presented with orientation patterns with step size 1°. Before present to the network, orientation patterns are corrupted with white noises of zero mean and 15° s.d.. Each orientation is presented for 5 s. The estimated orientations are decoded by population vector decoder from the responses of excitatory neurons, using a 200 ms moving window. Then, we computed the perceptual variability Var(*θ*) and bias (*θ*) for a given orientation *θ* by

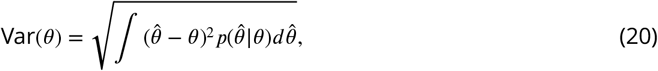

and

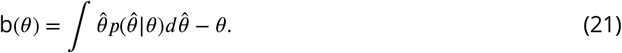

### Experiment details

#### Mouse strains

The mouse strain Pvalb-IRES-Cre (made by S Arbor at the Friedrich Miescher Institute for Biomedical Research, FMI) were obtained from the Jackson Laboratory and crossed with the reporter strain Rosa26-CAG-tdTomato (Ai9) (Jackson Laboratory) (***Chen et al., 2014***) to PV-Cre::Ai9 mice. The wild-type or transgenic mice were reared in the animal house on a 12/12 hr light/dark cycle for experiments. All animal procedures are contained in the animal experiment protocols, which have been reviewed and approved by the Animal Care Committees of the State Key Laboratory of Cognitive Neuroscience and Learning at Beijing Normal University (IACUC-BNU-NKLCNL-2016-02).

#### Visual stimulation

Visual stimuli were generated by our custom developed software using LabView (National Instruments) and MatLab (Mathworks), and were presented on a 20 inch cathode ray tube (CRT) monitor (Sony Multiscan G520; 30.5 x 30.5 cm; refresh rate, 60 Hz; maximum luminance, /), as described in our previous studies (***Chen et al., 2014, 2017*; *He et al., 2014***). The CRT monitor was placed 20 cm away in front of the mouse, subtending about 80° x 80° of the visual field. The full-screen drifting sinusoidal gratings of 12 different directions (spatial frequency, 0.02 Hz/degree; temporal frequency, 2 Hz) were used to measure the orientation tuned spike responses of spiking activity to stimuli in individual recorded neuron. Each stimulation trial consisted of 1 s of blank screen and 3 s of drifting gratings, and repeated for 4 times, in which stimulus orientations and directions were randomized.

#### Animal preparation for *in vivo* electrophysiology

Prior to electrophysiological recording, the animal was anesthetized by intraperitoneal injection of ketamine (50μ/g)/medetomidine (0.6μ/g) and mounted on a custom-built mouse stereotaxic device for recording in the visual circuit. The heart rate and body temperature of animals were monitored for the state of anesthesia, and additional half dose of anesthesia was given to sustain stable anesthesia if necessary during the surgery or recording. Body temperature was maintained at 37°C by a homeostatically controlled heating pad (RWD Life Science). Eye drops were applied when necessary to prevent the eyes from drying during the surgery or recording.

#### *in vivo* extracellular recording

Cell-attached recording on the layer 2/3 PV+ interneuon (with tdTomato red fluorescence) in the mouse V1 is achieved by the two-photon laser imaging guided target cell recording (***He et al., 2014***). A cranial window of 2 x 2 mm was made over the V1 area (***Madisen et al., 2010***). The dura mater in the window was carefully removed and a glass coverslip was mounted on the craniotomy using dental cement, allowing imaging fluorescent V1 cells to the depth of 150-μ (most within layer 2/3) *in vivo*. The targeted cell recording followed a procedure previously described in our study (4). Glass micropipettes filled with the artificial cerebrospinal fluid (aCSF) solution (mM: 124 NaCl, 2.5 KCl, 2 MgCl_2_, 2 CaCl_2_, 1.25 NaH_2_PO_4_, 26 NaHCO_3_ and 11 D–glucose (pH 7.35, B303 mOsm)) containing Alexa Flour 488 (50μ, Invitrogen) (tip opening, 2μ and 7– 10MΩ resistance) were advanced to the pia at a 14 degree angle with a micromanipulator (Sutter, MP-255). After contacting the pia, the pipette further advanced at the step of 1 mm toward tdTomato-expressing cells within the layer 2/3 under the guidance of continuous two-photon imaging. The 960-nm two-photon laser was used to excite Alexa 488 and tdTomato. As the pipette was advanced into the cortical tissue, a pressure of 0.2 psi was applied to the micropipette until the pipette touched the cell membrane, indicated by large drop in the magnitudes of currents responding to testing voltage pulse (5 mV, 100 Hz). The positive pressure was then released and a small amount of suction was immediately applied to form loose seal of resistances ranging from 30 to MΩ. This loose-seal configuration recorded spikes from a single cell without rupturing cell membranes. Spikes of PV+ cells were recorded.

The extracellular single-unit recording on the layer 2/3 PCs follows a procedure described in our previous studies (***Chen et al., 2014*; *He et al., 2014***), using the glass micropipettes filled with the aCSF (5– MΩ). Before the recording, a cranial window of 2 × 2 mm was made over the V1 area. In a penetration, only spiking activities with wide width were recorded from the neurons located in the corical depth less than μ (read by the micro-manipulator dial).

Electrical signals were recorded with an Axon MultiClamp 700B micropipette amplifier (Molecular Devices), filtered at 5 kHz (low pass), digitalized by a Digidata 1440A converter board (Molecular Devices) and finally acquired at 10 kHz with the pClamp10 (Molecular Devices) into a computer for further analysis. Spike events were detected and further analyzed with a custom program in the MatLab (Mathworks). The baseline spike activity was defined by the average spike number within 1 s before the onset of grating stimuli. Evoked spike rates in response to visual stimuli were measured over the duration of visual stimulation, with a subtraction from the baseline spike rate.

### Data analysis

#### Orientation selectivity

The level of orientation selectivity is quantified by global orientation selectivity index (1-CV):

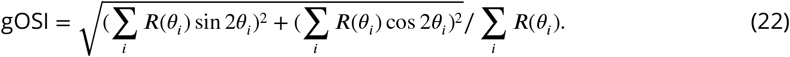

where *θ*_*i*_ is the orientation of the moving gratings, and *R*(*θ*_*i*_) is the spike rate (with baseline subtracted).

#### Preferred orientation

To determine the preferred orientation of a neuron, a Gaussian function was fitted to the mean spike rates evoked by drift gratings of 6 different orientations (see ***Figure 6***A, ***Figure 6***A), where the spike rate for every orientation was the average spike rate of the two corresponding opposite directions. Cells with *R*^2^ *>* 0.6 are included for further analysis. The preferred orientation was defined as the degree at which the fitted Gaussian function peaks.

#### Population tuning curve

To determine the sensory-evoked population activity, population tuning curves were computed as follows: First, we normalized the orientation tuning curve for each cell to its maximum spike rate (at the preferred orientation).Then, we averaged the normalized responses to different stimuli across neurons in the population.

#### Horizontal bias index

Horizontal bias index (HBI) was defined following (***Hagihara et al., 2015***): 1-dOri(preferredOrientation, horizontal)/45, which took a value between -1 (vertical) and 1 (horizontal), where dOri(X, Y) = min(|X-Y|, |180-|X-Y||) for each neuron.

## Acknowledgments

This work was supported by grants from the National Natural Science Foundation of China (32071025), the Beijing Municipal Science & Technology Commission (Z181100001518001) and the Interdisciplinary Research Fund of Beijing Normal University (to X-h.Z.). We thank Ye Li and Xiaolong Zou for helpful discussions and Wenhao Zhang for helpful advice on the draft of the paper.

---

**Appendix 0 Figure 9.**
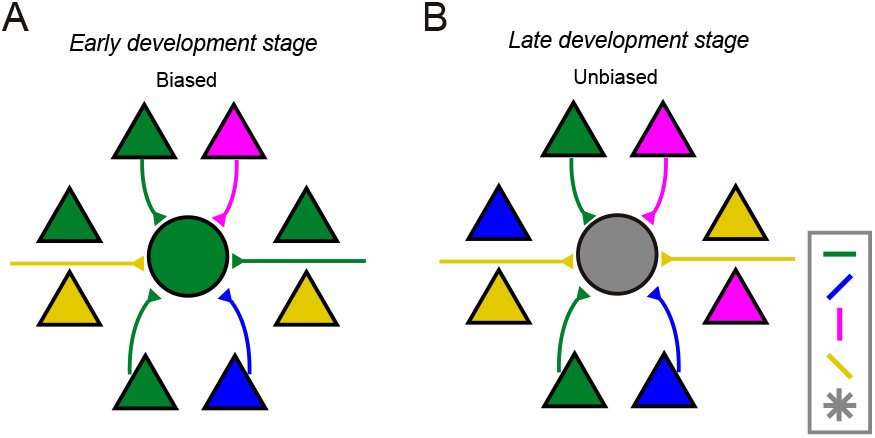
Developmental changes in orientation selectivity of PV+ interneurons reflects the developmental changes in the distribution of preferred orientations local PCs. **A**. Few days after EO, the local PC population is biased towards horizontal orientations, thus the PV neurons have high orientation selectivity and are likely to prefer horizontal orientations. **B**. Before CP, the horizontal bias in PC population is lost, and the number of PCs preferring to a specific orientation and its orthogonal orientations are likely to be equal, resulting in low orientation selectivity in PV+ inhibitory neurons

## Notes

https://github.com/AmazingAng/EmergeProbV1

### Competing Interest Statement

The authors have declared no competing interest.

### Summary of Updates

We change the color of the figure into CYMK format, which is friendly to color-blind readers.

